# EphA7 functions as a receptor for cell-to-cell transmission of Kaposi’s sarcoma-associated herpesvirus into BJAB B cells and for cell-free virus infection by the related rhesus monkey rhadinovirus

**DOI:** 10.1101/522243

**Authors:** Anna K. Großkopf, Sarah Schlagowski, Bojan F. Hörnich, Thomas Fricke, Ronald C. Desrosiers, Alexander S. Hahn

## Abstract

Kaposi’s sarcoma-associated herpesvirus (KSHV) is the causative agent of Kaposi’s sarcoma and is associated with two B cell malignancies, primary effusion lymphoma and the plasmablastic variant of multicentric Castleman’s disease. EphA2 functions as a cellular receptor for the gH/gL glycoprotein complex of KSHV for several adherent cell types. KSHV gH/gL was previously shown to also weakly interact with other members of the Eph family of receptors. Whether these interactions are of functional consequence remained unclear so far, even if other A-type Ephs were shown to be able to compensate for absence of EphA2 in overexpression systems. Here, we demonstrate for the first time that endogenously expressed EphA7 in BJAB B cells is critical for the cell-to-cell transmission of KSHV from producer iSLK cells to BJAB target cells. The BJAB lymphoblastoid cell line often serves as a model for B cell infection and expresses only low levels of all Eph family receptors other than EphA7. Endogenous EphA7 could be precipitated from the cellular lysate of BJAB cells using recombinant gH/gL, and knockout of EphA7 significantly reduced transmission of KSHV into BJAB target cells by over 80%. Finally, we demonstrate that receptor function of EphA7 is conserved between cell-to-cell transmission of KSHV and cell-free infection by the related rhesus monkey rhadinovirus (RRV), which is relatively even more dependent on EphA7 for infection of BJAB cells.

**IMPORTANCE:** Infection of B cells is biologically relevant for two KSHV-associated malignancies, the plasmablastic variant of multicentric Castleman’s disease (MCD) and primary effusion lymphoma (PEL). Elucidating the process of B cell infection is therefore important to understand the pathogenesis of these diseases. For various types of adherent cells, EphA2 has been shown by several groups to function as an entry receptor that is engaged by the gH/gL glycoprotein complex. Our previous findings indicate that KSHV does not only interact with EphA2, but can also bind to other members of the Eph family of receptor tyrosine kinases with comparatively lower avidity. We now analyzed the requirement of Eph interactions for infection of the BJAB B cell line, a model for infection of B cells by KSHV. We identify EphA7 as the principal Eph receptor for infection of this model B cell line by both KSHV and the related rhesus monkey rhadinovirus.

## INTRODUCTION

In addition to Kaposi’s sarcoma, Kaposi’s sarcoma-associated herpesvirus (KSHV) is associated with a variant of multicentric Castleman’s disease (MCD) and with primary effusion lymphoma (PEL) (1). Several publications demonstrate the importance of the cellular receptor EphA2 for KSHV entry into various adherent target cells (2–5). While the KSHV gH/gL complex exhibits the highest avidity for EphA2, it can also interact with other members of the Eph family of receptor tyrosine kinases (Ephs), similar to the gH/gL complex of the related rhesus monkey rhadinovirus (RRV) (6). However, so far it was unclear whether these interactions are of functional relevance for KSHV infection at endogenous protein levels. Interestingly, *in vitro* infection of established B cell lines by cell-free KSHV is extremely inefficient (7), while coculture of KSHV producing cells with target cells leads to robust infection (8). BJAB cells (9) are used as a model for B cell infection by KSHV (7, 8) and B cell biology in general (10). Our previous work had already demonstrated that cell-to-cell transmission of KSHV into BJAB cells is susceptible to inhibition of the interaction with receptors from the Eph family (6). An open question so far was which member of the Eph family of receptor tyrosine kinases is the principal receptor for infection of these cells. Additionally, we investigated the receptor requirements for BJAB infection by RRV, a closely related gamma2-herpesvirus of rhesus macaques (11) to further characterize its significance as an animal model virus for KSHV.

## RESULTS

We performed a two-step pulldown from the lysate of BJAB cells using Strep-tagged Fc fusion proteins of soluble versions of the gH proteins of KSHV, RRV 26-95 and RRV 17577 (12) in complex with the respective gL proteins as bait to identify cellular interaction partners (Figure 1A). While our group works with RRV isolate 26-95, we included isolate 17577 to identify possible differences in receptor interactions between the two described RRV isolates. Subsequent mass spectrometry analysis identified EphA7 as a prey in all three binding reactions (Figure 1B). In pulldowns with both RRV gH/gL complexes, EphA7 was the only membrane protein identified in excised bands in the 100-130kDa molecular weight range. In the KSHV pulldown we additionally found several peptides derived from EphA5. A comparison of the mRNA expression profiles of the 14 human Eph family receptors in a dataset (13) deposited in the Gene Expression Omnibus database revealed that BJAB cells predominantly express EphA7 (Figure 1C). We therefore focused our analysis on this member of the Eph receptor family. To confirm our mass spectrometry results, we repeated the pulldown with a similar experimental protocol and tested the precipitate for the presence of EphA7 by Western blot analysis. We could confirm that using soluble gH/gL complexes of KSHV and RRV 26-95 we pulled down a protein that reacted with an antibody to EphA7 from BJAB lysate, but not from 293T lysate (Figure 1D). In contrast, EphA2 was precipitated by KSHV gH/gL from 293T lysate but not from BJAB lysate, which mirrors expression of the two receptors in BJAB or 293T, respectively (Figure 1E). To test the functional relevance of the gH/gL-EphA7 interaction, we generated EPHA7 knockout (KO) cell pools by transducing BJAB with lentiCRISPRv2-based constructs targeting EPHA7. All four tested single guide RNAs (sgRNA) abrogated EphA7 expression compared to two non-targeting guide RNAs or two guide RNAs targeting EPHA2 as assayed by Western blot (Figure 2A). EphA2, the described KSHV receptor for adherent cells, is not expressed to detectable levels in BJAB cells (Figure 1E, 2B). The weak signal detected by the polyclonal EphA2 antibody in BJAB lysates appears at an apparent molecular weight below that of EphA2 in 293T cell lysate and is therefore most likely a non-specific reactivity, which would also be in accordance with mRNA expression profiles in databases and the absence of EphA2 in our initial pulldown experiment. After confirmation of the CRISPR/Cas9 knockout efficiency, we used the knockout cell pools to analyze receptor function of EphA7 for infection of BJAB cells. As BJAB cells are not readily amenable to infection with cell-free KSHV, we resorted to the previously described coculture cell-to-cell transmission system. This method allows for efficient infection of BJAB cells by overlaying chemically induced iSLK cells with BJAB cells and resolution of the two populations by flow cytometry after staining for expression of CD13 (as iSLK cell marker) and CD20 (as B cell marker) as described by Myoung et al. (8). In addition to iSLK cells harboring BAC16 KSHV wt we included iSLK cells harboring our previously described Eph-detargeted gH-ELAAN mutant in BJAB coculture experiments (Figure 2C) (14). When the results obtained with all sgRNA constructs targeting one specific gene were averaged (two non-targeting sgRNAs, two sgRNAs to EPHA2, four sgRNAS to EPHA7), knockout of EPHA7 resulted in a 86% reduction of infection with wt KSHV, whereas targeting EPHA2, which is not expressed at detectable levels as assayed by Western blot, resulted in a 27% reduction of wt KSHV infection, which was not significant. Analysis of all BJAB EPHA2^KO^ or EPHA7^KO^ cell pools compared to control cell pools in infection experiments with KSHV gH-ELAAN did not indicate significant changes between any of the groups. RRV 26-95, as opposed to KSHV, readily infects BJAB cells as free virus. Therefore, we infected the same set of knockout BJAB cells with cell-free wt RRV-YFP and with RRV-YFP gH-AELAAN, an Eph-detargeted RRV mutant, analogous to KSHV gH-ELAAN (Figure 2D). While the results obtained with RRV essentially paralleled those with KSHV, ablating EphA7 expression resulted in an even more pronounced and significant reduction in infection for all EPHA7-targeting constructs when compared to the non-targeting controls. When averaged, the BJAB EPHA7^KO^cell pools exhibited a 95% reduction in RRV-YFP infection vs non-targeting controls. Infection levels of RRV-YFP gH-AELAAN equaled RRV wt infection with matched genome copy numbers on BJAB EPHA7^KO^ cells, with no significant differences between controls and EPHA2^KO^ or EPHA7 ^KO^ cells.

**Figure 1.**
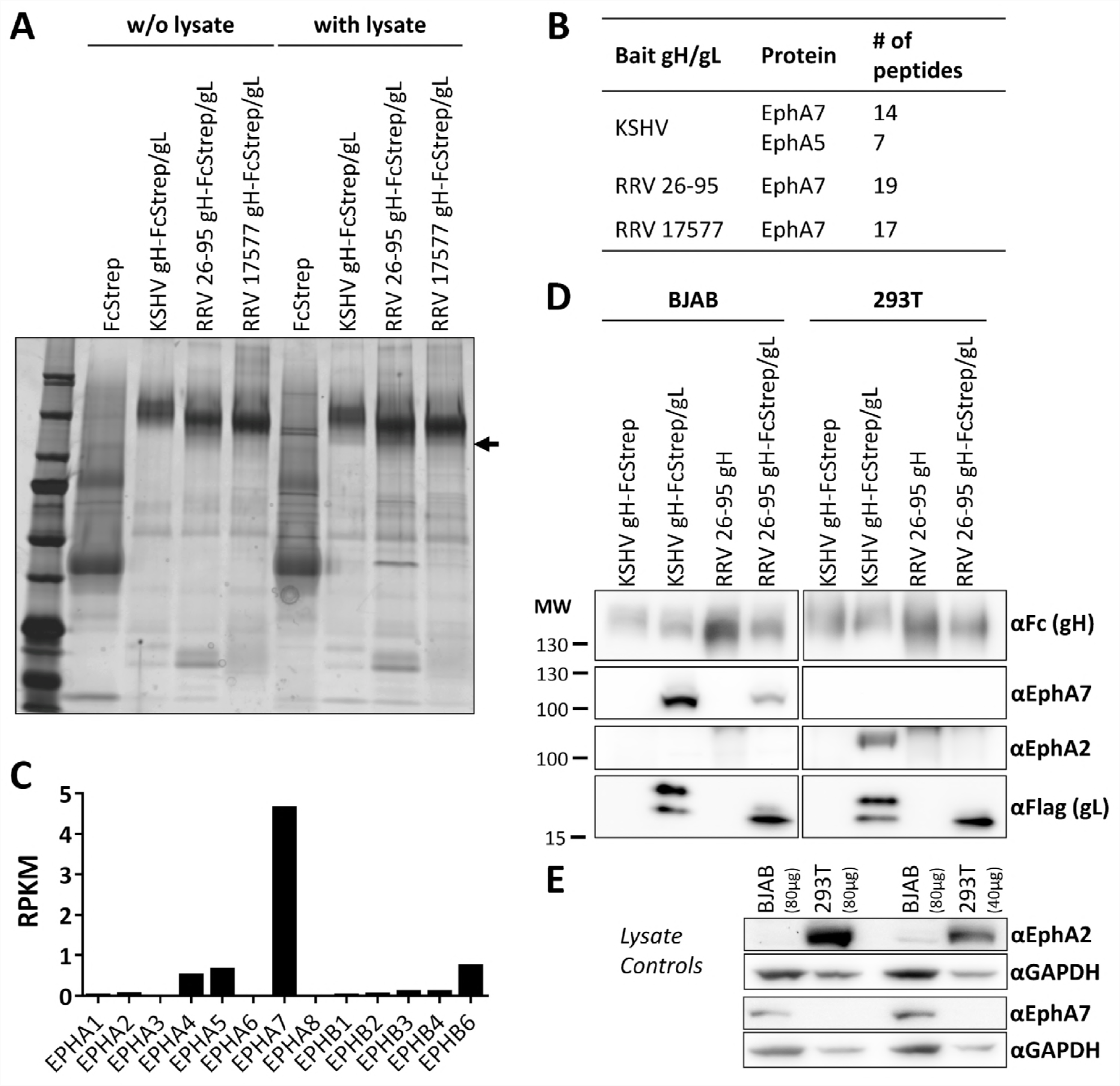
Identification of EphA7 as a gH/gL interacting protein in BJAB cells. A) Pulldown from BJAB cells with recombinant soluble gH-FcStrep/gL complexes. The precipitates were separated by PAGE, silver stained and bands at the indicated molecular weight (arrow) were excised and analyzed by mass spectrometry. B) Proteins and number of peptides per protein identified in each sample. C) RNA sequencing Reads Per Kilobase Million (RPKM) of the 14 EPH receptor genes as found in GEO dataset series GSE82184 BJAB-control (accession GSM2185732). D) Pulldown from BJAB and 293T lysate using recombinant soluble KSHV or RRV 26-95 gH/gL complexes as bait. Input amounts were normalized to wet cell pellet. Precipitates were analyzed by Western blot with the indicated antibodies. E) Expression control of EphA2 and EphA7 in BJAB and 293T cells. Lysates were analyzed by Western Blot. The BJAB:293T protein concentration ratio of 2:1 reflects the protein concentration ratio in the pulldown.

**Figure 2.**
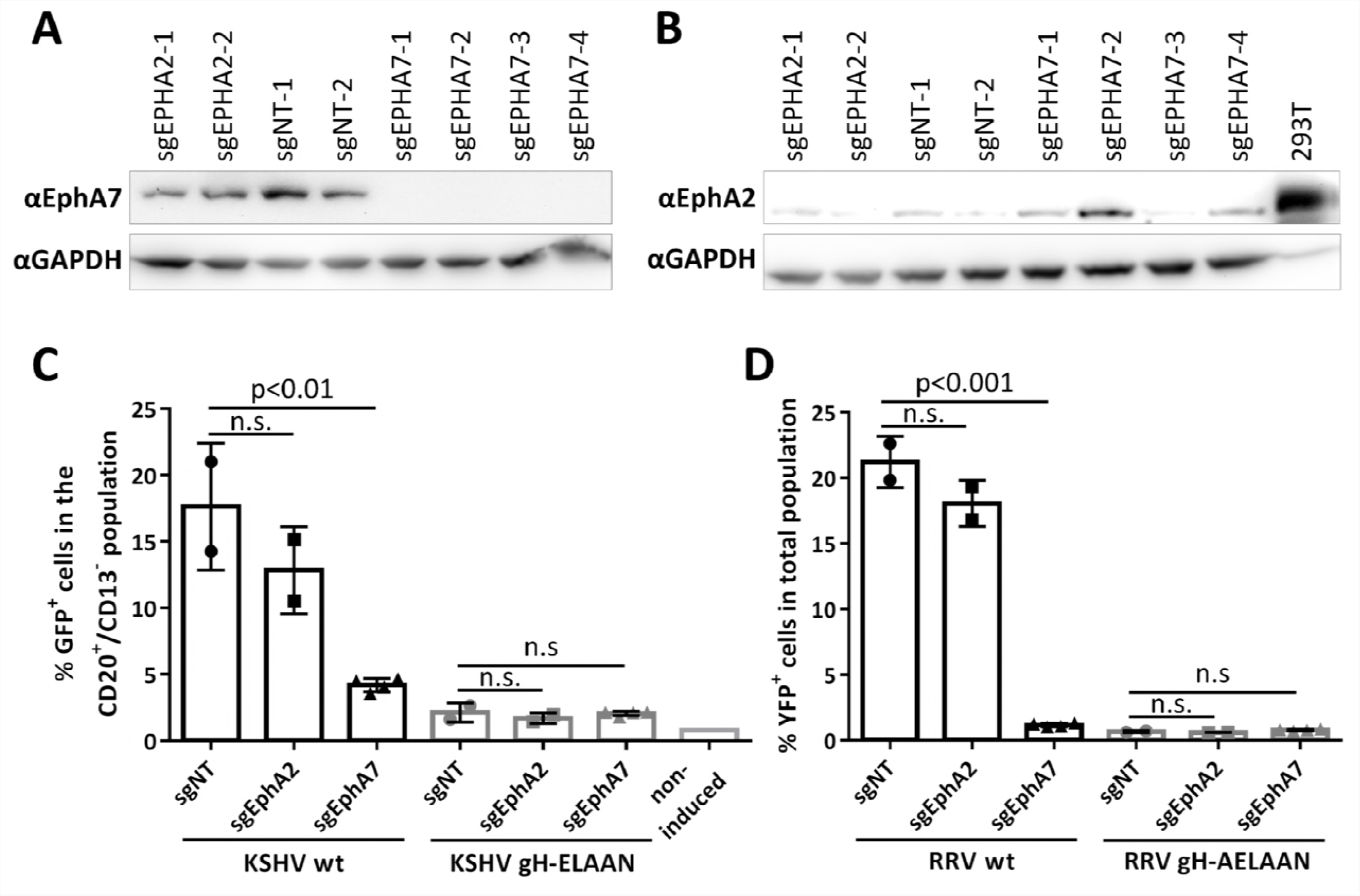
Knockout of EPHA7 reduces infection by KSHV and RRV. A) Knockout of EPHA7 in BJAB cells. BJAB cells were transduced with the indicated lentiCRISPRv2-based constructs, briefly selected, and cell pools were analyzed by Western blot. B) Absence of detectable EphA2 expression in BJAB cells. BJAB cells were treated as in (A) and analyzed for EphA2 expression by Western blot. One fourth of the protein amount loaded for the BJAB samples was loaded for the 293T EphA2 expression control. C) BJAB cells transduced with the lentiCRISPRv2-based constructs targeting the indicated genes were cocultured with iSLK cells harboring either BAC16 KSHV wt or BAC16 KSHV gH-ELAAN. Infection was analyzed by flow cytometry. The mean across groups of sgRNAs (n=3 infections per sgRNA) targeting EPHA2 (two sgRNAs), EPHA7 (four sgRNAs) or non-targeting (sgNT, two sgRNAs) is indicated by columns. The standard deviation of the mean is indicated by the error bars. The means of the individual triplicate infections for each sgRNA BJAB population within a group are given as symbols within the respective columns. ANOVA and pairwise comparison to the sgNT control group were used to test for significance. D) The same BJAB cell pools as in (C) were infected with cell-free RRV-YFP wt or RRV-YFP gH-AELAAN. Infection was analyzed by flow cytometry.

As we still detected residual infectivity upon EPHA7 knockout, we wanted to address the question, whether infection of EPHA7^KO^ BJAB cells could be further inhibited by a treatment with ephrinA4-Fc, a soluble Fc-fusion protein of a natural Eph ligand, which targets A-type Ephs and blocks infection of adherent cells by cell-free KSHV (6). This would indicate receptor functions of additional A-type Eph proteins other than EphA7. To this end we used a reduced set of our knockout cell pools and treated the cells either with PBS (control) or ephrinA4-Fc in PBS during the coculture experiment (Figure 3). While ephrin A4-Fc significantly inhibited infection of both EPHA2^KO^ BJAB cells and non-targeting control BJAB cells compared to PBS by over 85%, infection of EPHA7^KO^ BJAB cells was only reduced by 20% in the presence of ephrinA4-Fc, which did not reach significance. It should be noted that knockout of EPHA2 also lead to a small but significant reduction in infection in this experiment using only a reduced set of sgRNA constructs.

**Figure 3.**
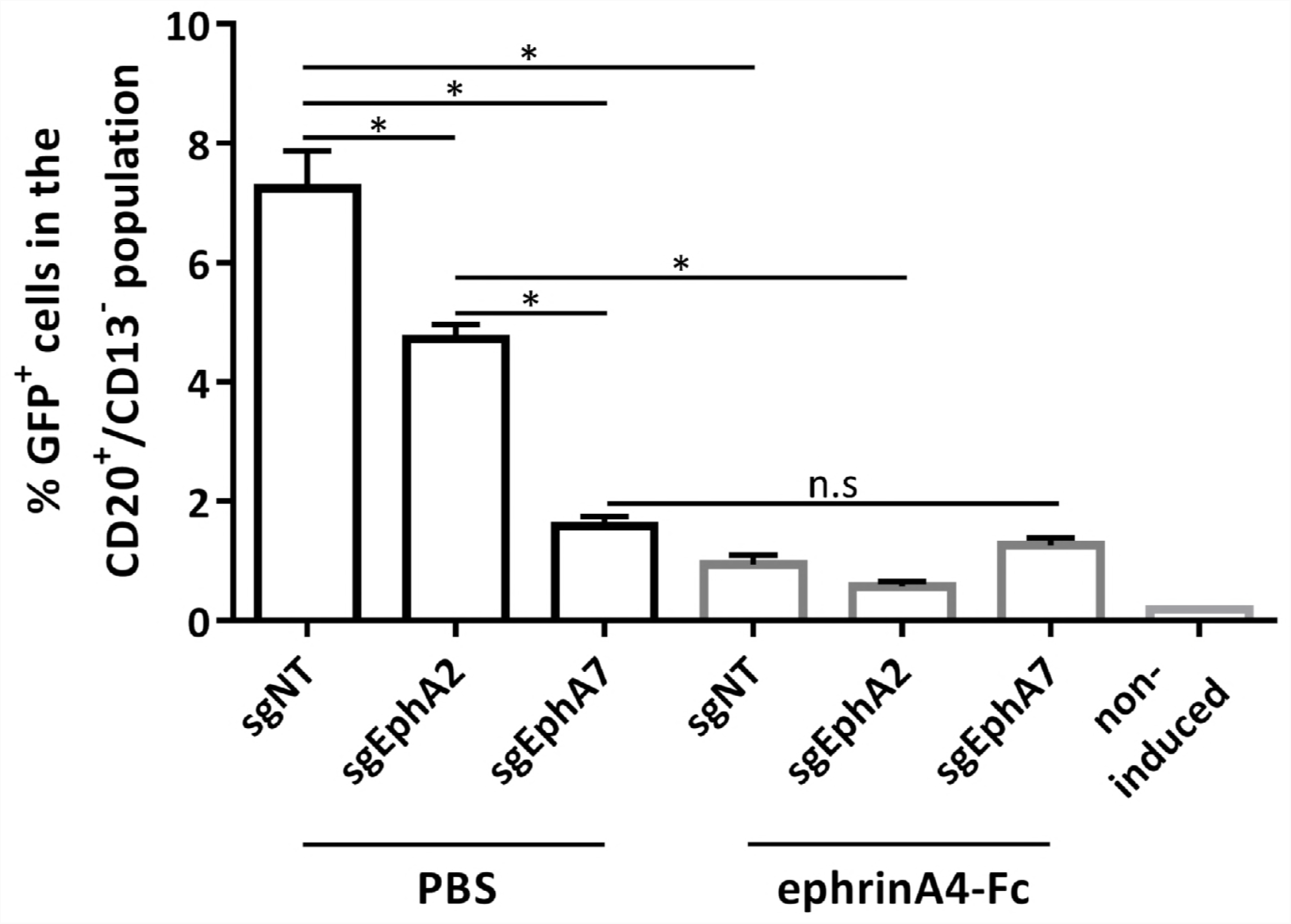
EphA7 is the predominant Eph receptor for infection of BJAB cells by KSHV. BJAB knockout cell pools (sgNT-1, sgEPHA2-1, sgEPHA7-1) were cocultured with iSLK cells harboring BAC16 KSHV wt. ephrinA4-Fc at a final concentration of 2µg/ml in PBS or PBS alone as control was added to the coculture. The indicated individual comparisons were made using ANOVA with Bonferroni correction. *: p-value < 0.05.

## DISCUSSION

We identified endogenous EphA7 in BJAB cell lysate as an interaction partner for the gH/gL complexes of both KSHV and RRV. This is not surprising insofar that it had already been shown in our previous publication that RRV binds relatively promiscuously to different Ephs, and that KSHV binds EphA2 with high affinity, but can also interact with other Ephs, among them EphA7, with lower affinity. We had also identified trace peptides of other Ephs alongside peptides of EphA2 after pulldown with KSHV gH/gL previously. Additionally, we had already identified EphA7 as an interaction partner for gH/gL of both RRV subtypes in 293T cells in a previous study (6). In contrast, we were not able to precipitate endogenous EphA7 from 293T cells with KSHV gH/gL under similar conditions, as was the case in previous studies. The reason for the lack of EphA7 – gH/gL interaction in 293T cells is most likely either competition by e.g. EphA2 for binding to the KSHV gH/gL complex and/or differences in relative expression of the different Ephs (Figure 1E). While a recent publication by TerBush et al. (2) has already suggested that Ephs other than EphA2 can function as receptors for KSHV, this has not been conclusively shown, as in this publication knockout of EPHA4 or EPHA5 e.g. in SLK cell did not lead to a reduction in KSHV infection. Quite to the contrary, EPHA4 knockout lead to an increase in infection. Demonstrating receptor function by a loss-of-function experiment such as gene knockout and not only after recombinant overexpression is therefore essential to demonstrate biological relevance at endogenous expression levels and in a physiological context. So far, only EphA2 has passed this test in several publications (2–5). We now show that knockout of EPHA7 in BJAB cells resulted in a significant reduction in infection by both KSHV and RRV. For RRV, the remaining level of infection was practically indistinguishable from that observed with the Eph-detargeted RRV-YFP gH-AELAAN mutant, indicating that EphA7 is the principal Eph receptor for infection of BJAB cells by RRV. For KSHV, EPHA7 knockout also reduced infection significantly by more than 85%. For both viruses, residual levels of infection with the Eph-detargeted virus mutants in both control and EPHA7^KO^ BJAB cells confirm our previous results that the interaction with Ephs contributes significantly to infection but is not strictly essential, most likely because alternative receptors exist. Unlike for RRV, the level of KSHV infection observed in the BJAB EPHA7^KO^ cultures was still marginally higher than that observed in the coculture experiments with iSLK harboring the Eph-detargeted KSHV gH-ELAAN. This difference in infection could be due to interaction with additional Eph receptors other than EphA7 by the KSHV gH/gL complex. The slight reduction of infection after transduction of sgRNAs directed against EPHA2, which reached significance in one set of infections (Figure 3) might hint at some residual function of EphA2 at very low expression levels. Our initial finding of some peptides derived from EphA5 in the analysis of the gH/gL pulldown as well as results by TerBush et al. demonstrating rescue of KSHV infection in EPHA2^KO^ SLK cells by recombinant EphA5 expression (2) furthermore suggest that EphA5 may also have a function in mediating the remaining level of infection after knockout of EPHA7. However, we only observed a very slight reduction of KSHV infection by treating the iSLK BAC16 KSHV wt/ BJAB EPHA7 ^KO^ coculture with ephrinA4-Fc, which did not reach significance. In the light of this inability to block KSHV infection of EPHA7 ^KO^ cells with ephrinA4-Fc and given the very small difference in infection between KSHV wt on EPHA7 ^KO^ cells and infection with the Eph-detargeted KSHV gH-ELAAN overall, any attempts of analyzing the function of additional receptor interactions in this context would not be informative. Nevertheless, we think that it is highly likely that other Ephs, which like EphA5 weakly but detectably interact with KSHV gH/gL (6), may play a role similar to EphA7 for KSHV infection of other cell types, in particular in the absence of the high-affinity EphA2 receptor. This should be addressed in separate studies using loss-of-function methodology. Overall, while KSHV exhibits the highest affinity for EphA2, our results demonstrate that KSHV uses other Ephs under circumstances where these are more abundantly expressed, which parallels the interaction of ephrins with their receptors (15) and receptor usage by RRV (6). Whether there is a direct correlation between affinity and receptor function or whether the mechanism is more complicated will also be a subject for future studies. A recent study on the role of glycoprotein K8.1 A in the infection of B cells, but not in the infection of other cell types,hints at possible mechanistic differences (16) specific for B cell infection, although infection of BJAB by cell-to-cell transmission was not analyzed in that study. Another open question is whether the function of EphA7 in cell-to-cell transmission is somewhat different from the function of e.g. EphA2 in the infection of epithelial or endothelial cells, and whether the specific nature of cell-to-cell transmission may allow for more efficient use of receptors with comparatively lower affinity for gH/gL.

## METHODS AND MATERIALS

### Cells and virus

BJAB cell lines were obtained from the Leibniz-Institute DSMZ-Deutsche Sammlung von Mikroorganismen und Zellkulturen GmbH. iSLK cells were a kind gift from Jinjong Myoung (17), 293T cells from Stefan Pöhlmann, and primary rhesus monkey fibroblasts from Rüdiger Behr. BJAB cells were propagated in RPMI medium (Thermo Fisher Scientific) supplemented with 10% fetal bovine serum (FBS) (Thermo Fisher Scientific) and 50µg/ml gentamycin (PAN Biotech). 293T cells were cultured in Dulbecco’s Modified Eagle Medium (DMEM), high glucose, GlutaMAX, 25mM HEPES (Thermo Fisher Scientific) supplemented with 10% FBS and 50μg/ml gentamycin. iSLK cells were maintained in DMEM supplemented with 10% FBS, 50μg/ml gentamycin, 2.5μg/ml puromycin (InvivoGen) and 250μg/ml G418 (Carl Roth). RRV-YFP, RRV-YFP gH-AELAAN and iSLK cells harboring BAC16 KSHV wt or BAC16 KSHV gH-ELAAN were produced as described previously (14).

### Coculture infections

iSLK cells were seeded in 48well plates at 100 000 cells per well. After five hours, lytic replication was induced with 1µg/ml doxycycline (Sigma) and 2.5mM sodium butyrate (Carl Roth) over night in DMEM, high glucose, GlutaMAX, 25mM HEPES with 10% FBS. After one day, medium was exchanged to 400µl RPMI with 10% FBS and 50µg/ml gentamycin and the respective BJAB cell pools at 40 000 cells per well. For blocking experiments, recombinant ephrinA4-Fc protein (R&D Systems) was added at the start of the coculture to a final concentration of 2µg/ml.

### Flow cytometry analysis

Cocultured cells were harvested by pipetting and fixed with 2% formaldehyde (Carl Roth) in PBS. The cells were stained with anti-CD13 (clone WM15, PE coupled, BIOLEGEND) and anti-CD20 (clone 2H7, Alexa647 coupled, BIOLEGEND) antibodies at a 1:50 dilution in 5% FBS in PBS for 30min – 45min. After 2 washes in PBS, the cells were post-fixated in 2% formaldehyde in PBS. The samples were analyzed on a LSRII flow cytometer (BD Biosciences). Flow cytometry data was further analyzed using Flowing software (Version 2.5), for details see supplemental figure S1. Statistical analysis was carried out using GraphPad Prism version 6 for Windows (GraphPad Software, La Jolla California USA).

### Generation of knockout cell pools

Knockout cell pools were generated using the lentiCRISPRv2 system (18) as described previously (19), with the exception that transfection was carried out using PEI (20). In short, BAJB cells were transduced with lentiviruses harboring the indicated sgRNAs. After 48h the selection antibiotic puromycin (Invivogen) was added to a final concentration of 10µg/ml. After initial selection the puromycin concentration was reduced to 1µg/ml. The following single guide RNAs (sgRNAs) were inserted into plentiCRISPRv2 (a kind gift from Feng Zhang (Addgene plasmid number 52961)): 2 non-targeting controls sgNT-1 (plasmid Ax127): ATCGTTTCCGCTTAACGGCG, sgNT-2 (Ax128): TTCGCACGATTGCACCTTGG, 4 sgRNAs directed against EPHA7, namely sgEPHA7-1 (Ax279): GGAGAATGGTTAGTGCCCAT, sgEPHA7-2 (Ax280): GACATGTGTCAGCAGTGCAG, sgEPHA7-3 (Ax281): GGATTTCCTCTCCACCCAAT, sgEPHA7-4 (Ax282):

GATTTCCTCTCCACCCAATG, 2 sgRNAs directed against EPHA2, namely sgEPHA2-1 (Ax122): CTACAATGTGCGCCGCACCG, sgEPHA2-2 (Ax123): GGACTTTGCTGCAGCTGGAG. sgRNA sequences for all sgRNAs directed against EphA7 as well as EPHA2-2 were determined using E-CRISP (21). The lentiCRISPRv2-sgEPHA2-1 plasmid was purchased from Genscript. NT sgRNA sequences were taken from the GeCKO (version 2) library (18).

### Pulldown experiments and Western blot

Two-step pulldown with specific elution and re-precipitation using gH-FcStrep/gL complexes for identification by mass spectrometry was performed as described previously (8) with 1ml of wet BJAB cell pellet in 4ml of lysis buffer as starting material for pulldown per sample. The samples were then separated by polyacrylamide gel electrophoresis using 8-16% gradient gels (Invitrogen) and the gel was silver stained using the SilverQuest staining kit (Life Technologies). Mass spectrometry analysis of individual gel bands was carried out by the Taplin mass spectrometry core facility, Harvard Medical School. For pulldown followed by Western blot analysis, cells were lysed with 1ml of lysis buffer (1% NP40 150mM NaCl 1mM EDTA 25mM HEPES pH7.3 with addition of protease inhibitor cocktail (Amresco)) per ml of wet cell pellet. The lysate was clarified by centrifugation 21 100g for 20min and reacted with gH-FcStrep/gL-Flag complexes that were pre-coupled to Strep-Tactin XT (IBA) beads. After three washes with lysis buffer, the precipitates were analyzed by polyacrylamide gel electrophoresis and Western blot as described previously (14) using antibodies to EphA7 (clone E-7, sc-393973) and EphA2 (C-20, sc-924), both from Santa Cruz Biotechnology, 1:100 and 1:500, respectively, in NET gelatine (150mM NaCl, 5mM EDTA 50mM Tris, 0.05% Triton-X-100, 0.25% gelatin, pH 7.5) and donkey anti-mouse HRP-coupled (Dianova) or goat anti-rabbit HRP coupled (Life Technologies) secondary antibody in 5% dry milk powder in PBS with 0.05% Tween20. Membranes were imaged using Luminata forte substrate (Merck) on an INTAS ECL ChemoCam system. Similar lysis conditions were used for analysis of gene knockout cell pools by Western blot.

## ACKNOWLEDGEMENTS

This work was supported by grants to A.S.H. by the Deutsche Forschungsgemeinschaft (HA 6013/1, HA 6013/4-1), by grant RO1 AI072004 from the National Institutes of Health (NIH) to R.C.D., and by base grant RR00168 from NIH to the New England Primate Research Center. We thank Rüdiger Behr for primary rhesus monkey fibroblasts, Stefan Pöhlmann and Jens Gruber for reagents.

## SUPPLEMENTAL FIGURE LEGEND

Suppl. Figure S1. Gating strategy for scoring of KSHV infection in iSLK/BJAB cocultures. The percentage of GFP^+^ cells in the CD13^-^CD20^+^ population is given for controls and one representative iSLK BAC16 KSHV wt/BJAB control or iSLK BAC16 KSHV wt/BJAB EPHA7^KO^ sample. Numbers in brackets represent event counts (CD13^-^CD20^+^ GFP^+^ events/ CD13^-^CD20^+^ events). It should be noted that the ‘iSLK BAC16^induced^ w/o BJAB’ control sample, which represents the assay background from e.g. spill-over, contained one CD13^-^CD20^+^ GFP^+^ ‘false positive’ background event in 10 000 total events. This represents no more than 1.54% of the CD13^-^CD20^+^ GFP^+^ population in any sample.

